# The non-adrenergic imidazoline-1 receptor protein Nischarin is a key regulator of astrocyte glutamate uptake

**DOI:** 10.1101/2020.11.28.402222

**Authors:** Swati Gupta, Narges Bazargani, James Drew, Souvik Modi, Hélène Marie, David Attwell, Josef T. Kittler

## Abstract

Astrocytic GLT-1 is the main glutamate transporter involved in glutamate buffering in the brain, pivotal for glutamate removal at excitatory synapses to terminate neurotransmission and for preventing excitotoxicity. We show here that the surface expression and function of GLT-1 can be rapidly modulated through the interaction of its N-terminus with the nonadrenergic imidazoline-1 receptor protein, Nischarin. The phox domain of Nischarin is critical for interaction and internalization of surface GLT-1. Using live super-resolution imaging, we found that glutamate accelerated Nischarin-GLT-1 internalization into endosomal structures. The surface GLT-1 level increased in Nischarin knockout astrocytes, and this correlated with a significant increase in transporter uptake current. Furthermore, Nischarin knockout in astrocytes is neuroprotective against glutamate excitotoxicity. These data provide new molecular insights into regulation of GLT-1 surface level and function and suggest novel drug targets for the treatment of neurological disorders.

**Highlights:**

- The phox domain of Nischarin interacts with the N-terminal tail of the main astrocyte glutamate transporter, GLT-1.
- Nischarin promotes internalization of GLT-1 to endosomes.
- Glutamate modulates GLT-1 surface levels via regulation of the Nischarin-GLT-1 interaction.
- Genetic loss of Nischarin significantly increases GLT-1 surface expression, resulting in increased glutamate transport currents and enhanced neuroprotection.

## Introduction

Glutamate transport into cells is mediated by excitatory amino acid transporters (EAATs). Of the five EAAT subtypes found in the CNS, EAAT2/GLT-1 is predominantly expressed in astrocytes and is a major means for glutamate clearance from the extracellular space (Danbolt, 2001). Glutamate buffering through transporter binding helps to maintain a low extracellular glutamate concentration that facilitates termination of fast excitatory synaptic transmission (Diamond and Jahr, 1997, Wadiche et al., 1995a, Wadiche et al., 1995b, Tong and Jahr, 1994). Moreover, lateral diffusion of surface GLT-1 can also regulate glutamate clearance and thus shape glutamatergic neurotransmission (Al Awabdh et al., 2016, Murphy-Royal et al., 2015). The transporters have a long transport cycle (~70 msec) (Wadiche et al., 1995a) compared to the time scale of glutamate presence at the synapse (~1 msec) (Barbour and Hausser, 1997). However, despite this long transport cycle, efficient glutamate clearance occurs during synaptic activity due to the high surface density of transporters (Danbolt, 2001). A low extracellular glutamate concentration (below the submicromolar level that tonically activates NMDA receptors) is also crucial to prevent excitotoxic cell death (Choi et al., 1987, Choi, 1987). Thus, control of GLT-1 density on the astrocyte surface via molecular mechanisms modulating its intracellular trafficking is crucial for normal synaptic physiology and prevention of excitotoxicity.

Four isoforms of GLT-1 have been identified, which exhibit similar functional properties and oligomerize to form homomeric and heteromeric GLT-1 pools. However, the GLT-1 isoforms differ in their N- and C-termini, allowing for interaction with different intracellular proteins and offering an opportunity for differential regulation of the isoforms during physiological and pathological conditions (Peacey et al., 2009). So far, scaffolding proteins, such as PSD-95, PICK-1 and MAGI-1 have been shown to interact with the PDZ domain containing C-terminus of the GLT-1b isoform (Underhill et al., 2015, Sogaard et al., 2013, Zou et al., 2011, Gonzalez-Gonzalez et al., 2008). Here, we have identified the non-adrenergic imidazoline-1 receptor protein Nischarin, as a novel physiological GLT-1 N-terminus interacting protein in astrocytes. Nischarin is a cytoplasmic protein that shows a diverse set of functions, including regulation of the cytoskeletal network (through Rac1), interaction with endosomes (via its phox domain), and regulation of receptor (mu opioid and integrin α_5_ receptors) surface levels (Alahari et al., 2000, Keller et al., 2017, Li et al., 2016, Kuijl et al., 2013, Alahari et al., 2004, Alahari, 2003, Dong et al., 2017).

Here, we report that the Nischarin phox domain is sufficient for interaction with a 30 amino acid stretch of the GLT-1 N-terminus. Super-resolution imaging revealed that glutamate drives internalization of GLT-1 in a Nischarin-dependent pathway. In addition, Nischarin knockout (Nisch^KO^) animals exhibit increased astrocytic GLT-1 surface expression that correlates with increased transporter current and enhanced neuroprotection. Together, these data suggest a novel mechanism for how glutamatergic signalling is regulated in the central nervous system under both physiological and pathological conditions.

## Results

### GLT-1 and Nischarin interact *in vivo*

A yeast two-hybrid screen (Y2H) with the GLT-1 N-terminus identified a clone that encoded the phox domain of the Nischarin protein as a novel interactor for the GLT-1 protein (Marie et al., 2002). Next, a coimmunoprecipitation assay was performed in COS cells co-transfected with GLT-1a (tagged in the extracellular loop with a V5 epitope) and GFP-tagged Nischarin (Nisch) lacking the phox domain (GFP-NischΔphox), or Nischarin’s phox domain alone (GFP-phox) or a GFP control (Fig. 1A). We found that the GFP-phox domain construct, but not the GFP-NischΔphox, co-precipitated with GLT-1 (Fig. 1A), suggesting that the phox domain of Nischarin is sufficient for the interaction with GLT-1. In parallel experiments, co-immunoprecipitation assays in brain lysates generated from adult wild-type (WT) and GLT-1 knockout (GLT^KO^) animals confirmed the interaction between Nischarin and GLT-1 protein in WT mice, which was lacking in the GLT^KO^ mice (Fig. 1B). Western blotting in cortical astrocytes derived from rat pups (P0-P2) confirmed expression of endogenous Nischarin (Suppl. Fig.1A).

**Figure 1:**
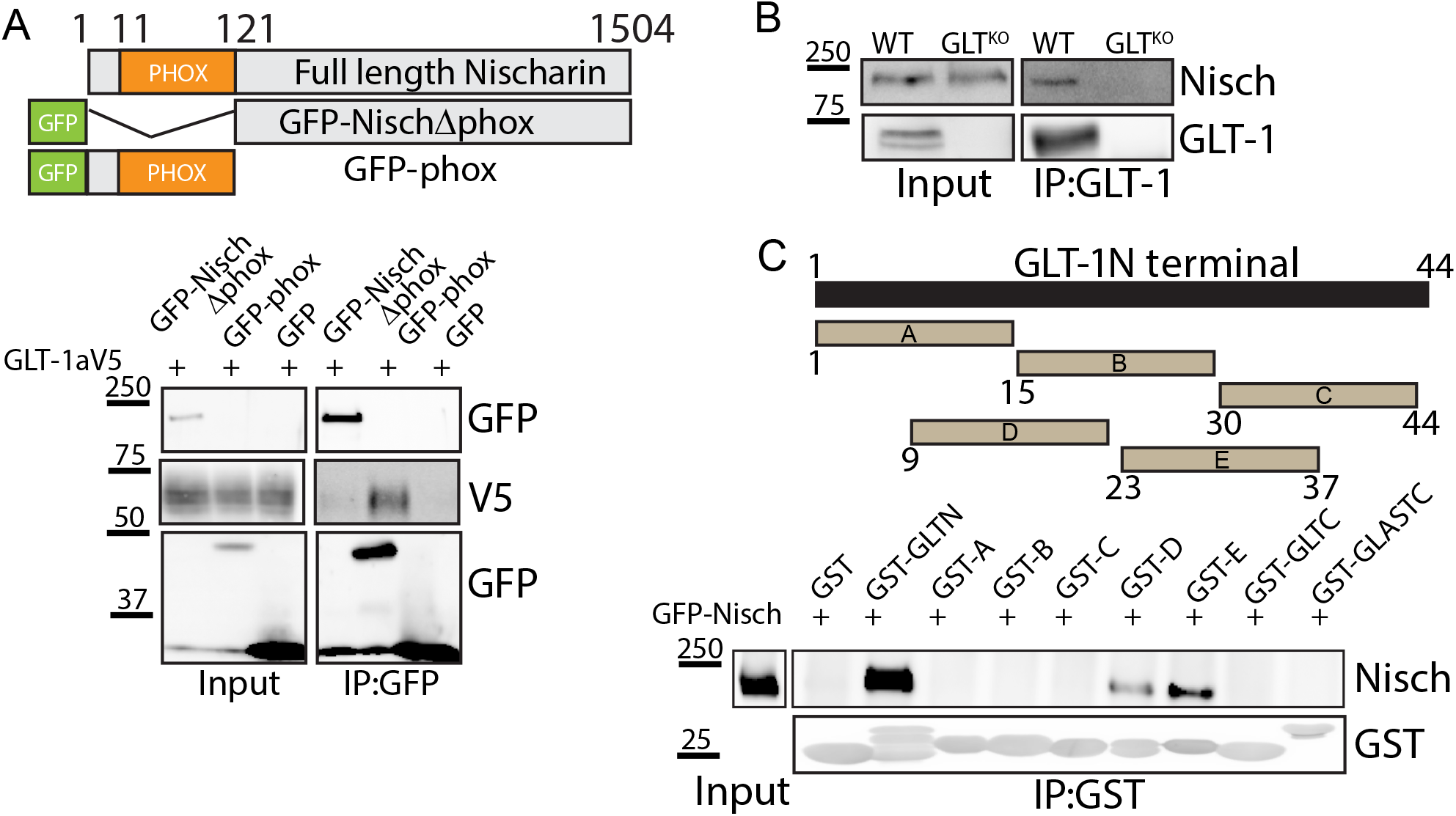
The phox domain of Nischarin interacts with GLT-1. **A)** Schematic diagram depicting full length Nischarin and GFP-tagged mutants. GLT-1aV5 coimmunoprecipitated with GFP-phox and not GFP-NischΔphox mutant or GFP control. **B)** Coimmunoprecipitation experiments from mouse brain homogenate from WT and GLT^KO^ mice, showing Nischarin to be part of a native complex with GLT-1 (n=3). **C)** Schematic diagram of GST fusion constructs for GLT-1 N-terminus, 15 amino acid stretches of GLT-1 N-terminus (A-E), GLT-1 C-terminus and GLAST C-terminus. GFP-Nisch was successfully pulled down with full length GST fused GLT-1 N-terminus and to GST fusions D (amino acids 9-23) and E (amino acids 23-37).

Using a GST fusion assay, we further narrowed down the binding region within the N-terminal tail of GLT-1 participating in the interaction with Nischarin. The following GST tagged fusion proteins were used as bait; 1) full length GLT-1 N-terminus fusion protein, 2) fusion proteins containing overlapping stretches (15 amino acid in length) of the GLT-1 N-terminus (A-E), 3) full length GLT-C-terminus, and 4) full length GLAST C-terminus, to assess pull down of Nischarin from lysates of GFP-Nisch or GFP transfected COS cells. While the N-terminus of GLT-1 successfully interacted with Nischarin, its C-terminal end and that of GLAST did not.

Interestingly, within the GLT N-terminal, two 15 amino acid stretches from 9-24 and 23-37 successfully interacted with Nischarin (Fig. 1C). Given that the other GST fused GLT-1 N terminal segments did not interact with Nischarin, this implies that the Nischarin binding site on GLT-1 is complex and that the two sites (9-24 and 23-27) are sufficient for binding Nischarin, independently.

### Nischarin promotes endocytosis and not recycling of GLT-1

Nischarin has been shown to alter the surface levels of receptors, including integrins and mu opioid receptors (Li et al., 2019, Li et al., 2016, Lim and Hong, 2004). To determine whether Nischarin regulated GLT-1 surface density, we used an ‘antibody feeding’ immunofluorescence internalization assay in HeLa cells to visualize GLT-1 trafficking. HeLa cells were co-transfected with GLT-1 tagged in its extracellular domain with HA (GLT-1a-HA) along with either GFP-Nisch or GFP as a control. Briefly, antibody against the HA tag, present in the extracellular loop of GLT-1, was incubated with the cells for 15 min prior to placing them in the incubator at 37°C for 60 min to allow for internalization, followed by differential immunostaining of the surface and internal GLT-1 pool. To obtain a reference value for basal surface GLT-1 levels prior to internalization, cells were fixed immediately after the antibody surface labelling (constituting the T_0min_ population). At baseline (T_0min_), in control cells, robust surface and low internal GLT-1 labeling was observed. Even after 60 min, a significant change in GLT-1 internalization was not observed in the control (Fig. 2A, 2C), suggesting stable turnover of the transporter under basal culture conditions. In GFP-Nisch expressing cells, although no significant difference was observed in GLT-1 distribution initially at T_0min_, by 60 min significant increases were observed in the accumulation of GLT-1 within intracellular compartments. (Fig. 2B, 2C). This suggests that Nischarin regulates constitutive trafficking of GLT-1 under basal conditions.

**Figure 2:**
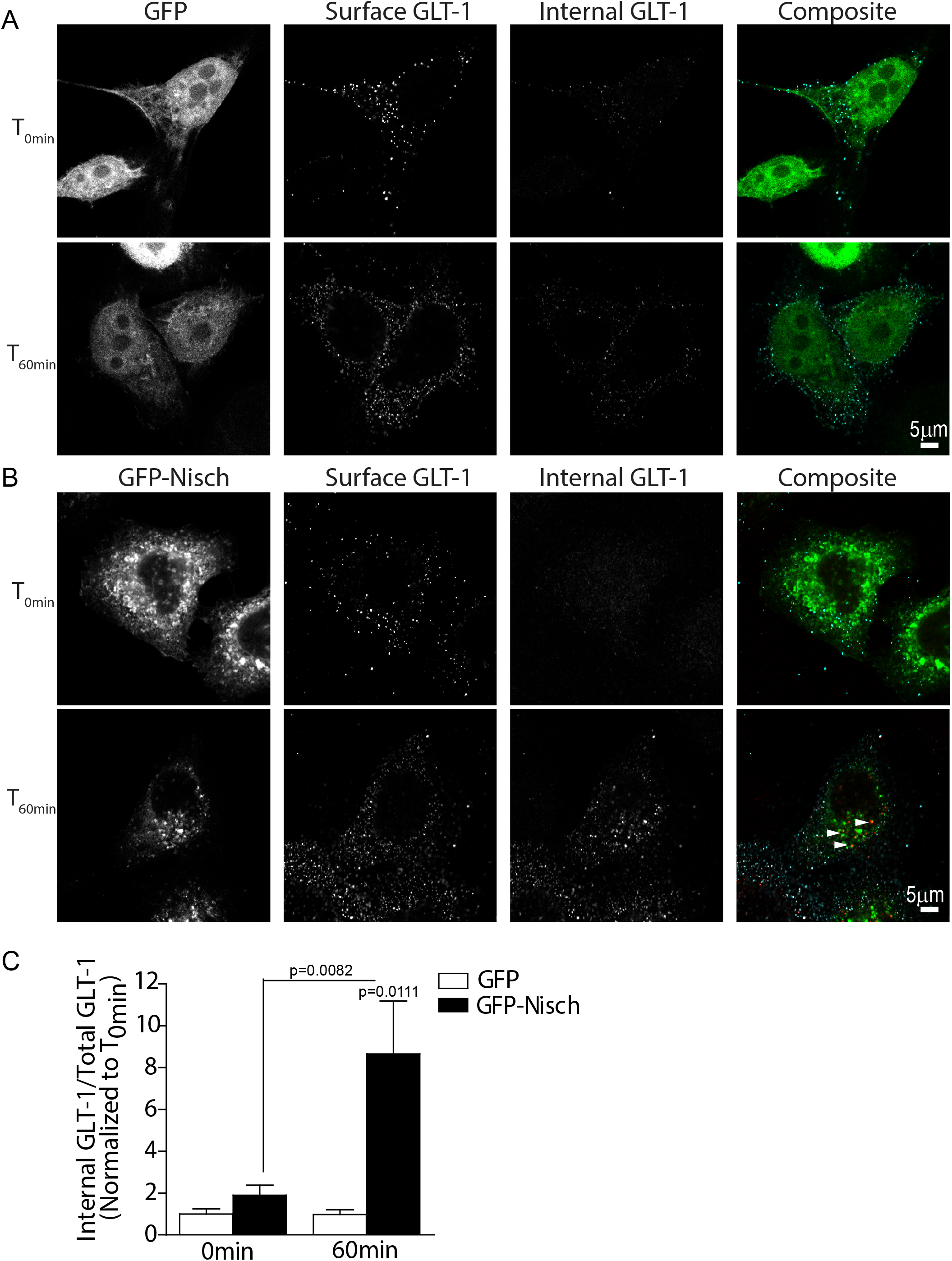
Antibody feeding assay revealed that Nischarin promotes internalization of GLT-1. Surface and internal GLT-1 populations labelled in HeLa cells co-expressing **A**) GFP and GLT-1a-HA or **B**) GFP-Nisch and GLT-1a-HA at 0min and 60min. **C**) Quantification of internalized GLT-1 to total GLT-1 levels at T_60 min_ relative to T_0 min_. One-way ANOVA, Kruskal-Wallis test with Dunn’s correction (*n* = 12).

As trafficking of GLT-1 is dependent on endocytic and recycling pathways, we next assessed the effects of Nischarin on GLT-1 recycling. No significant differences were found between the GLT-1 recycling rates in GFP-Nisch cells and control (Supp. Fig. 1B-D). Together, these data suggest that Nischarin promotes translocation of GLT-1 from the surface to intracellular compartments, but (in contrast to its effect on mu opioid receptors: Li et al., 2016) does not affect GLT-1 recycling.

Next, astrocytes were transfected with either GFP-Nisch or GFP-NischΔphox or GFP control, and co-cultured with hippocampal neurons. At DIV 14, immunostaining studies revealed significant co-localization between GFP-Nisch positive endosomal structures and the early endosomal marker EEA1. The GFP-Nisch positive vesicles showed variability in size and shape and were distributed throughout the astrocyte cell body and processes. However, GFP-Δphox overexpressing cells showed a cytosolic expression of GFP-Δphox, corroborating previous reports that Nischarin is targeted to endosomes via its phox domain (Suppl Fig. 2A-D). Furthermore, endogenous GLT-1 co-localized significantly with intracellular vesicles positive for GFP-Nisch (Fig 2E). Together these data suggest that intracellular GLT-1 accumulates in Nischarin positive early endosomal structures within the astrocyte cell body and processes.

### Glutamate promotes Nischarin mediated GLT-1 intracellular trafficking in fixed and live hippocampal astrocyte cultures

It has been previously reported that glutamate treatment decreases clustering and surface expression of GLT-1 (Al Awabdh et al., 2016, Underhill et al., 2015). The PDZ binding domain containing protein DLG1 interacts with the C-terminal PDZ ligand of GLT-1b to regulate its surface density (Underhill et al., 2015). However, the role of molecules interacting with the N terminus of GLT-1, remains under explored. We therefore investigated whether Nischarin regulates GLT-1 trafficking under activity driven conditions, which we mimicked by applying glutamate.

Using a surface biotinylation assay, the effect of glutamate on surface GLT-1 levels in Nischarin overexpressing astrocytes was assessed. Pure cortical astrocyte cultures were co-transfected with GFP and GLT-1a tagged with V5 (control) or GFP-Nisch and GLT-1a-V5. The transfected cultures were either left untreated or exposed to glutamate (100 μM, 1 h). Glutamate treatment significantly decreased surface GLT-1 level compared to the untreated control (Fig. 3A), as expected (Ibanez et al., 2016). Nischarin overexpression significantly decreased the GLT-1 surface levels compared to the untreated control in the absence of glutamate, corroborating our findings in Fig. 2. Glutamate treatment in Nischarin over-expressing astrocytes did not cause a further decrease in surface GLT-1 levels in comparison to Nischarin overexpression or glutamate application alone, suggesting that overexpression of Nischarin alone is sufficient to drive the internalisation of surface GLT-1.

**Figure 3:**
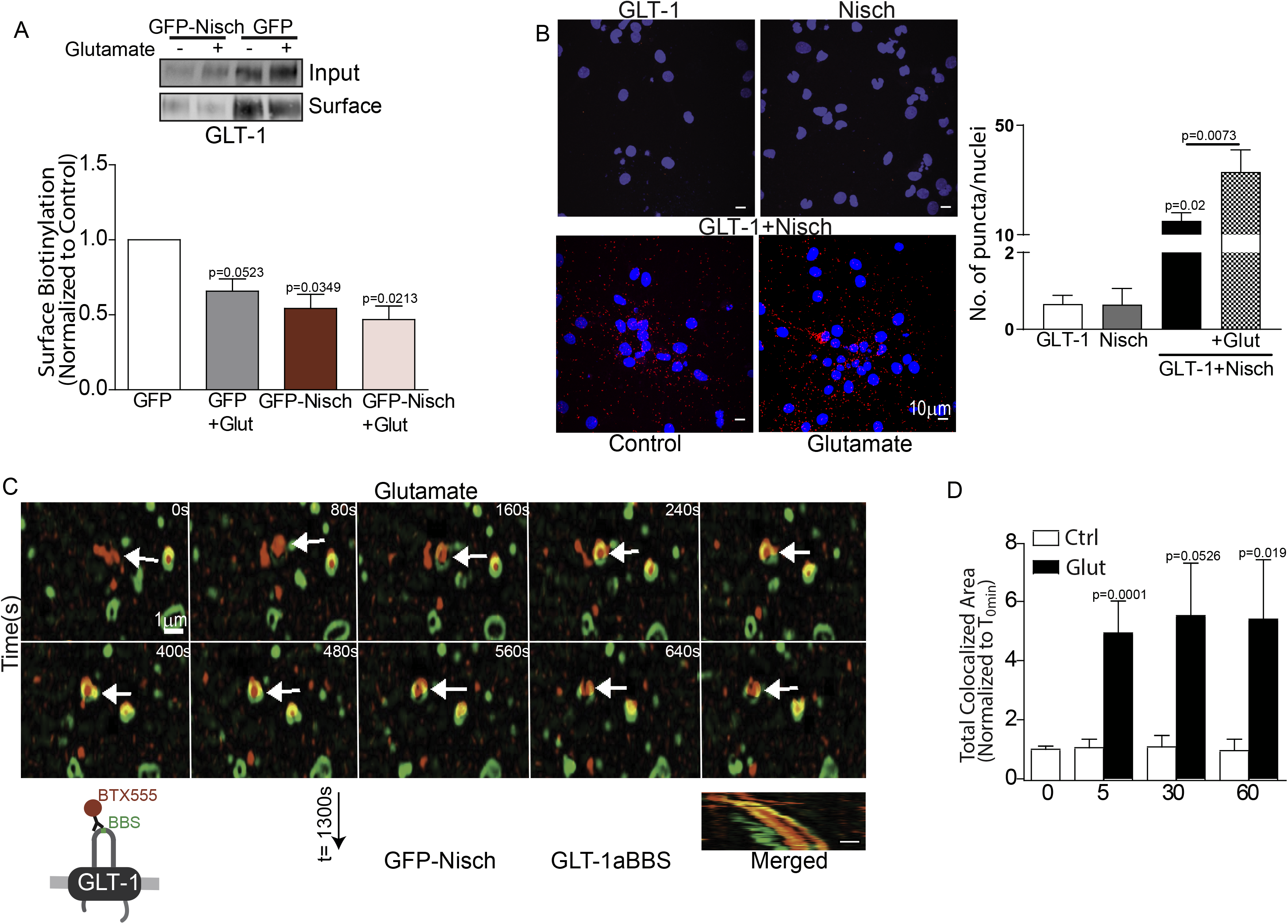
Nischarin mediates glutamate-dependent GLT-1 internalization in astrocytes. **A)** Surface biotinylation assay showing surface GLT-1 level in astrocytes transfected with GFP and GLT-1a-V5 or GFP-Nisch and GLT-1a-V5 following +/− 100μM glutamate treatment. One-way ANOVA, post hoc Dunnett’s multiple comparison test (*n* = 4 individual experiments). **B)** Proximity ligation assay in DIV14 hippocampal culture. Increased red puncta per nuclei (DAPI stained (blue)) is indicative of increased direct interaction between Nischarin and GLT-1 in hippocampal culture. Glutamate treatment (100μM, 1h) significantly increased GLT-1-Nischarin interaction compared to control. One-way ANOVA, post hoc Tukey’s test (n = 3 individual preparations). **C)** Schematic representation of GLT-1BBS construct bound to BTX conjugated Alexa-555 (BTX555). Astrocytes expressing GFP-Nisch and GLT-1aBBS were labelled using BTX555 and dual colour live structured illumination microscopy monitored trafficking of GLT-1 following glutamate treatment. Merged kymographs of GFP-Nisch vesicle (green) and GLT-1 bound BTX-555 (red) reveal co-localized diagonal trajectory, representing moving vesicles. **D)** Quantification of GFP-Nisch and GLT-1aBBS expressing astrocytes treated with 100μM glutamate for 0, 5, 30 and 60min showed increased co-localization between Nisch and GLT-1 compared to untreated controls. P values by unpaired t-test, Mann Whitney test (*n* = 6-14).

Next, using a proximity ligation assay (PLA), a powerful tool that detects a positive protein interaction only if the two proteins are closer than 40 nm, we determined the effect of glutamate on the endogenous GLT-1-Nischarin interaction. In DIV14 hippocampal cultures, under control conditions, a significant increase in Nischarin-GLT-1 association (assessed by number of red puncta per DAPI labelled (blue) nuclei) was observed compared to single antibody (Nisch/GLT-1) controls. Upon glutamate treatment (100 μM, 1 h), the number of puncta observed were significantly increased compared to the control (Fig. 3B). Together with the biotinylation assay, these results suggest that glutamate enhances the endogenous GLT-1-Nischarin interaction, and that this drives internalization of surface GLT-1.

To monitor live trafficking of GLT-1, we took advantage of a high affinity 13 amino acid α-bungarotoxin (BTX)-binding site (BBS) that has been exploited for tracking AMPA receptor movements in and out of the cell membrane (Sekine-Aizawa and Huganir, 2004). We explored a similar strategy to assess the time course of glutamate’s action on the GLT-1-Nischarin interaction, by engineering the extracellular loop of the GLT-1 transporter (Fig. 3C) to include the BBS tag. This position in GLT-1 transporters is silent in terms of its impact on receptor structure and function (Peacey et al., 2009). The astrocytes were co-transfected with GLT-1a-BBS and GFP-Nisch and co-cultured with hippocampal neurons. The hippocampal co-culture was incubated with Alexa-555-conjugated BTX (BTX555) at 37°C for 20 min to allow live labelling of surface GLT-1. Live time lapse imaging using structured-illumination microscopy (SIM) tracked transporter internalization and individual endosomal events in labelled GLT-1a-BBS astrocytes (co-cultured with hippocampal neurons) co-expressing GFP-Nisch and exposed to ACSF alone or ACSF containing 100μM glutamate (Fig. 3C). GFP-Nisch positive intracellular vesicles were observed within the astrocyte processes and cell body. Glutamate application resulted in internalization of BTX555 labelled surface GLT-1 transporters to GFP-Nisch labeled vesicles, as seen by the diagonal and colocalized lines in the kymographs representing GLT-1 and Nischarin vesicle movements within the astrocyte process (Fig. 3C). GFP-Nisch was found to be associated with the inner plasma membrane, and the internalized surface GLT-1 was trafficked into GFP-Nisch-positive vesicles, pinched off from the plasma membrane (Suppl. Video 1, 2).

Using the same setup as described above, the co-cultures were exposed to varying durations (5, 30, or 60 min) of glutamate (100 or 0 μM) and subsequently fixed and imaged using confocal microscopy to ascertain the time course of Nischarin mediated GLT-1 trafficking. GLT-1 showed increased colocalization with GFP-Nisch labeled vesicles (in white, Suppl. Fig. 3A) upon glutamate treatment in comparison to control at all 3 time points (Fig. 3D). Together, these results reinforce the role of Nischarin in regulating GLT-1 internalization during glutamate application, with colocalization observed as early as 5 min.

### GLT-1 transporter density and function are altered in Nisch^KO^ mice

We further characterized the role of Nischarin mediated regulation of astrocytic GLT-1 by using transgenic Nisch^KO^ mice (Suppl. Fig. 3B-D). Lack of Nischarin expression was confirmed in homozygous Nisch^KO^ mice, whereas the heterozygous (Nisch^HET^) mice showed reduced Nischarin protein expression in comparison to the WT control (Fig. 4A). A surface biotinylation assay revealed significant increases in surface GLT-1 levels in astrocytes derived from Nisch^KO^ transgenic mice compared to from WT control mice. The surface GLT-1 level in astrocytes derived from Nisch^HET^ transgenics were not significantly different from that of WT (Fig. 4A). Immunostaining studies also revealed a significant increase in total astrocytic GLT-1 mean intensity in hippocampal co-cultures derived from Nisch^KO^ mice in comparison to WT control (Fig. 4B). These data support Nischarin’s role in regulating GLT-1 surface levels.

**Figure 4:**
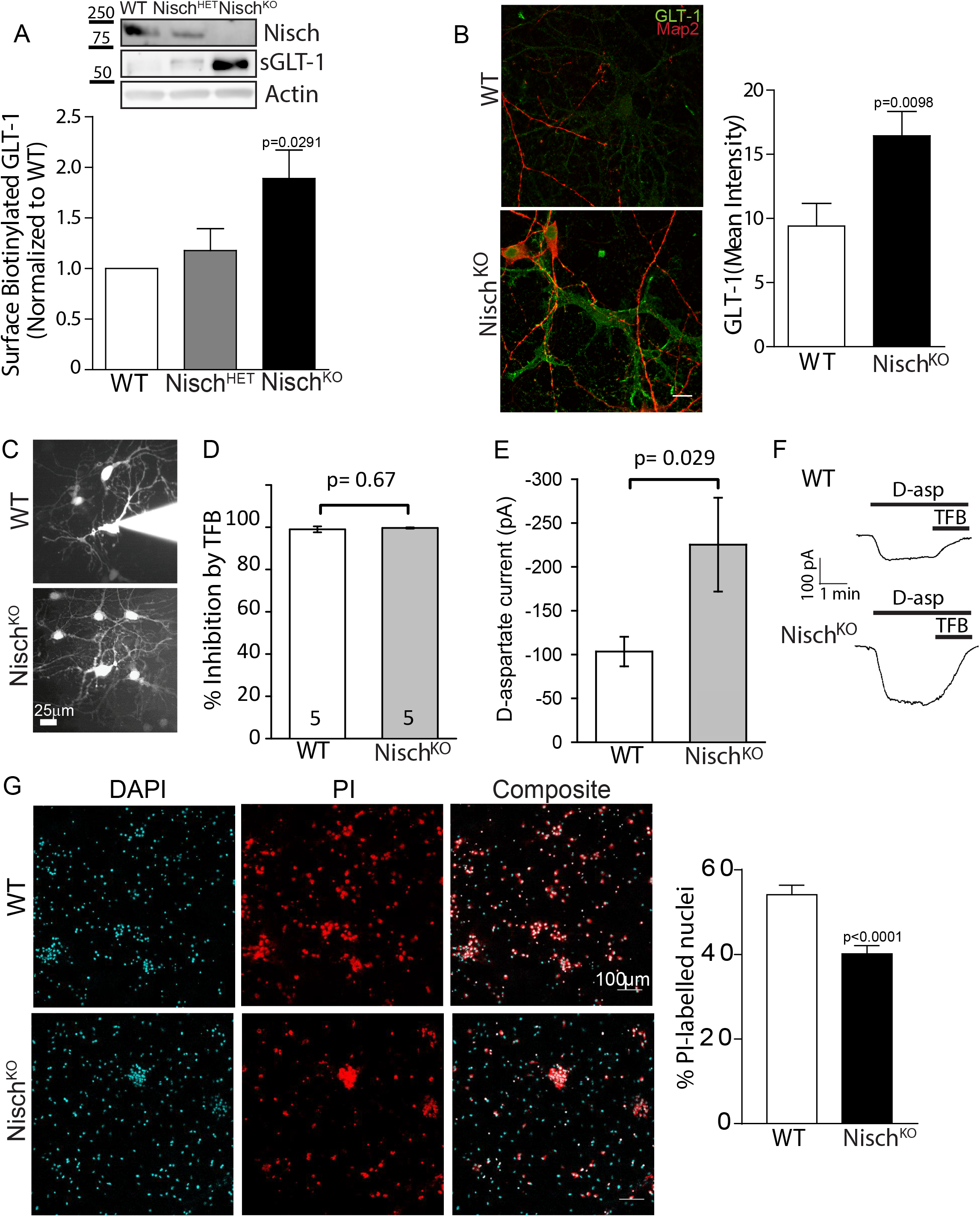
GLT-1 surface density and transporter uptake current are enhanced in Nisch^KO^ astrocytes. **A)** Western blot analysis in cortical astrocytes derived from WT, Nisch^HET^ and Nisch^KO^ E16 embryos, confirmed decrease and loss of Nischarin in the HET and KO cultures. Surface biotinylation assay showed significant increase in GLT-1 surface density in KO culture compared to WT control. One-way ANOVA, post hoc Tukey’s test (*n* = 3 animals). **B)** Representative images for GLT-1 (green) and Map2 (red) immunostaining in astrocytes derived from DIV14 WT and Nisch^KO^ hippocampal culture. A significant increase in GLT-1 mean fluorescence intensity was observed in Nisch^KO^ astrocytes. Unpaired Student t-test (*n* = 11-13). **C)** Examples of astrocytes filled with Alexa 488 from the patch pipette (still attached to the cell for the WT and after electrode removal for the Nisch^KO^) in WT and Nisch^KO^ hippocampal tissue cultures. **D)** The D-aspartate evoked current is completely blocked by TFB-TBOA. **E)** A significantly larger D-aspartate evoked current was recorded in the Nisch^KO^ astrocytes compared to the WT, unpaired student t-tests. **F)** Sample traces showing the D-aspartate evoked (200 μM) current and its inhibition by the GLT-1 and GLAST transporter blocker TFB-TBOA (TFB, 10 μM), in WT and Nisch^KO^ astrocytes. **G)** Representative confocal images showing nuclear staining DAPI (cyan) and PI labelling (red) in DIV 14 WT and Nisch^KO^ hippocampal culture, 24h following a glutamate insult. Bar graph showing percentage of PI-labelled nuclei (*n* = 3, unpaired two-tailed t-test).

Given the increased surface GLT-1 level in Nisch^KO^ astrocytes, we undertook functional studies where glutamate uptake was assessed using whole cell patch-clamp of astrocytes in hippocampal neuron-glial co-cultures (Brew and Attwell, 1987). For each glutamate anion transported into astrocytes by GLT-1, three Na^+^ and one H^+^ are also transported in, and one K^+^ is exported from the cell (Levy et al., 1998a, Zerangue and Kavanaugh, 1996, Levy et al., 1998b). Thus, two net positive charges are imported per glutamate taken up, and therefore uptake can be measured from the current it produces. Astrocytes were whole cell patch clamped and their identity was confirmed by the following: 1) dye filling (Alexa 488 or 594) to show coupling with other astrocytes (Fig. 4C), 2) their low input resistance (Suppl. Fig. 4A) and 3) their negative resting potential (Suppl. Fig. 4B). The input resistance and resting potential were not significantly affected by knock-out of Nischarin (Suppl. Fig. 4A, B). After applying blockers of action potentials (150 nM TTX), inward rectifying K^+^ channels i.e. the main conductance of astrocytes (200 μM BaCl_2_), and glutamate and GABA receptors (blocked with 10 μM NBQX, 50 μM D-AP5, 10 μM 5,7-dichlorokynurenate, 10 μM MK-801 and 10 μM bicuculline), once a steady membrane current was reached, at a voltage near the cell’s resting potential (−90 mV), glutamate transporters were activated by applying D-aspartate (200 μM). D-aspartate evoked an inward current in both WT and Nisch^KO^ astrocytes (Fig. 4E). The currents were confirmed to be mediated by glutamate transporters, since they were blocked by the GLT-1 and GLAST blocker TFB-TBOA (10 μM Fig. 4D, F). Consistent with the surface biotinylation assay, we found that the glutamate uptake current was almost two-fold higher in the Nisch^KO^ astrocytes in comparison to the WT astrocytes (p=0.029, Fig. 4E, F).

Dysfunction of glutamate clearance can cause overstimulation of glutamate receptors and result in neuronal injury, termed excitotoxicity. To further investigate the neuroprotective function of astrocytes, we carried out an excitotoxicity assay using hippocampal neuron-astrocyte co-cultures derived from WT and Nisch^KO^ mice (at DIV 14). The cultures were challenged with 10 μM glutamate and 10 μM glycine for 24 hours. Neuronal death was analyzed using propidium iodide (PI) and DAPI staining, and the number of PI-positive nuclei (red) were counted. Glutamate treatment evoked significantly less PI-labelling of neurons in Nisch^KO^ co-cultures compared with neurons in wild type co-cultures (Fig. 4G). These data suggest that when challenged with neurotoxic glutamate levels, lack of Nischarin is protective against cell death, presumably due to the increased glutamate uptake that lack of Nischarin results in.

## Discussion

Glutamatergic neurons are responsible for majority of the excitatory synaptic transmission and plasticity occurring in the brain. The astrocytic glutamate transporters serve the critical role of efficiently clearing glutamate from the extracellular space (Tanaka et al., 1997) to ensure normal glutamate signalling. GLT-1 is one of the highest expressed proteins in the brain (1% of total brain protein: (Lehre and Danbolt, 1998). The high number of surface glutamate transporters (GLT-1 at a density of 8,500 transporters/μm2 as well as GLAST at 2,500 transporters/μm2) compensate for the slow transport cycle (12-70ms) to ensure effective clearance of the ~4000 glutamate molecules released from a single synaptic vesicle (Lehre and Danbolt, 1998, Murphy-Royal et al., 2017). GLT-1 undergoes activity-dependent surface diffusion, and glutamate-bound GLT-1 from ‘synapse facing’ sites are continuously replaced with GLT-1 lacking bound glutamate to help maintain a high concentration of available surface transporters at the astrocytic plasma membrane (Murphy-Royal et al., 2017). Thus, an in-depth understanding of the molecular mechanisms regulating and maintaining the surface GLT-1 density is crucial. Here, we have identified a novel interacting protein partner, Nischarin, a non-adrenergic imidazoline-1 receptor (Alahari, 2003) that regulates intracellular trafficking of GLT-1 in response to the neurotransmitter glutamate.

We found that Nischarin interacts with the N-terminal tail of GLT-1 through its phox domain. Specifically, Nischarin co-precipitated with amino acids 9-37 within the intracellular, unstructured N-terminal tail of GLT-1. Amino acids 9-23 have also been implicated in the interaction with Ajuba, a scaffolding protein that allows GLT-1 to regulate intracellular signalling or interact with the cytoskeleton (Marie et al., 2002). These amino acids are conserved across the four GLT-1 isoforms (Peacey et al., 2009), suggesting Nischarin could regulate all four isoforms, unlike previously identified regulators of GLT-1 trafficking that bind specifically to the PDZ domain found in GLT-1b (Bassan et al., 2008, Underhill et al., 2015). Additionally, a co-immunoprecipitation assay using brain lysates confirmed that the Nischarin-GLT-1 interaction occurs in intact brain. The findings reported here raise the intriguing possibility that, in addition to Nischarin’s previously reported role in regulating cytoskeletal signalling, cell migration, Rab-dependent endosomal sorting, and regulation of integrins and mu opioid receptors (Keller et al., 2017, Kuijl et al., 2013, Alahari, 2003), it may have an activity-dependent role in modulating glutamate concentration at the synapse.

GLT-1 is known to undergo constitutive and regulated endocytosis, which determines its availability for glutamate clearance from extracellular compartments in the nervous system (Martinez-Villarreal et al., 2012). Our antibody feeding assays have revealed that overexpression of Nischarin redistributed surface GLT-1 transporters into endosomal structures, but did not alter transporter recycling under basal conditions. Taken together, our data not only support an interaction between Nischarin and GLT-1 but also indicate the possibility that Nischarin alters trafficking of GLT-1 by sequestration. The phox domain of Nischarin is a stretch of ∼110 amino acids that is a phosphatidylinositol 3-phosphate-binding (PI3P) module, and PI3P is enriched in early endosomal membranes (Lim and Hong, 2004). Our data confirmed previous findings that Nischarin is targeted to early endosomes, marked by EEA1 (Kuijl et al., 2013), but also showed multiple Nischarin positive yet EEA1 negative vesicular structures, revealing a much wider distribution of Nischarin within the endosomal system. Furthermore, we observed that the GLT-1-Nischarin vesicular structures are spatially distributed along the astrocyte cell body and processes.

Surface biotinylation studies revealed that astrocytic GLT-1 surface levels decreased (40%) in response to glutamate (Fig. 3A), which is consistent with previous findings (Ibanez et al., 2016). Overexpressing Nischarin in astrocytes exposed to glutamate treatment did not significantly further alter surface GLT-1 levels, implying that Nischarin occludes the effect of glutamate on surface GLT-1. Alteration in surface GLT-1 levels by Nischarin could offer a means for modulating glutamatergic activity at the synapse. Given the increased Nisch-GLT-1 interaction following glutamate exposure, a likely explanation for these results is that Nischarin mediates the effects of glutamate dependent GLT-1 surface density regulation. The recruitment of Nischarin to GLT-1 could have additional consequences as Nischarin is known to act as a scaffolding platform for signalling pathways through its interactions with multiple proteins including, integrin α_5_, PAK, Rac, LIMK and ERK (Alahari, 2003, Alahari et al., 2000, Juliano et al., 2004). Further work is also needed to establish whether the blood pressure lowering effects of imidazoline drugs such as clonidine are in any way mediated by effects on glutamate transport.

Using time-lapse monitoring of the transporter (employing a GLT-1aBBS construct that can bind fluorophore conjugated BTX, eliminating the use of bulky antibodies which could promote clustering and affect membrane trafficking properties (Sekine-Aizawa and Huganir, 2004) we showed that glutamate binding and/or transport triggered intracellular trafficking of GLT-1 into Nischarin labelled intracellular compartments within 5 min of glutamate exposure. This time course suggests that the GLT-1-Nisch trafficking could be of more relevance in pathological conditions such as ischemia or traumatic brain injury, where the extracellular concentration of glutamate remains elevated (in the 100-200 μM range) for hours (Ibanez et al., 2016). SIM resolution allowed tracking of single GFP-Nisch labelled vesicles containing GLT-1BBS bound BTX555, and the resultant kymograph confirmed co-localization and inwardly directed (towards the cell body), slow (~minutes) movement as vesicles traversed the astrocytic process.

Astrocytes derived from Nisch^KO^ transgenic animals exhibit increased surface GLT-1 density and a concomitant two-fold increase in transporter uptake currents. This enhanced surface GLT-1 density served to reduce cell death after glutamate insult, demonstrating the relevance of this mechanism to pathology. Dysregulation of Nischarin regulation of GLT-1 transporter surface density and function could affect glutamate clearance. Ineffective glutamate clearance is observed in many neurodegenerative diseases, including amyotrophic lateral sclerosis, epilepsy, Alzheimer’s, Huntington’s and Parkinson’s disease (Hindeya Gebreyesus and Gebrehiwot Gebremichael, 2020, Peterson and Binder, 2019). Together, this work not only reveals a novel mechanism by which GLT-1 intracellular trafficking and function are regulated but also provides possible new avenues of research for treating neurological disorders.

## Acknowledgements

This work was supported by grants from the BBSRC (BB/I00274X/1) and European Research Council grant 282430 (Fuelling Synapses) to J.T.K. S.M. was supported by an EMBO Long-Term Fellowship and Marie Skłodowska-Curie International Incoming Fellowship (Nos. 630033 and 913033). This work was also supported by a PhD studentship from the Medical Research Council (MRC) to J.D. (1477260) and a Wellcome Trust 4 year PhD studentship in Neuroscience to N.B. D.A. was supported by an Investigator Award (099222/Z/12/Z) from the Wellcome Trust. We thank all members of the Kittler laboratory for helpful discussions and comments. We also thank the Light Microscopy facility at the MRC Laboratory for Molecular Cell Biology SURF facility for training and technical support.

## Figure Legends

**Suppl. Movies 1 and 2.**: Single internalization events showing inward movement of GFP-Nisch positive intracellular vesicle (green) and GLT-1-BBS-BTX555 (red) captured using time lapse confocal microscopy under basal conditions and following glutamate treatment (100 μM). Images were acquired every 20 sec; movie accelerated to 10 fps. Scale bar 5 μm.

**Suppl. Figure 1:**

**A**) Representative western blots for endogenous Nischarin present in cortical neuron culture and pure astrocytic culture.

Nischarin does not affect GLT-1 recycling. HeLa cells co-expressing **B**) GFP and GLT-1a-HA or **C**) GFP-Nisch and GLT-1a-HA were live-labelled with anti-HA and rate of recycling was assayed using antibody feeding. **D**) No difference in recycled GLT-1 levels was observed in the two groups at T30 min and T_60 min_. At T_60 min_, both GFP control and GFP-Nisch overexpressing cells showed significant recycling of GLT-1 to the surface compared to T0 min, One-way ANOVA, post hoc Tukey’s test (*n* = 12).

**Suppl. Figure 2:** Structured illumination microscopy showing **A**) GFP-Nisch or **B**) GFP-NischΔphox or **C**) GFP and endogenous GLT-1 (red) and EEA1 (magenta) in astrocytes of DIV14 hippocampal culture.

**D**) Significant colocalization was observed between GFP-Nisch and the endogenous endosomal marker, EEA1 in comparison to the GFP-NischΔphox mutant and GFP control. One-way ANOVA, Kruskal Wallis test with Dunn’s multiple comparison test (*n* = 18-24 cells).

**E**) Significant colocalization was observed between GFP-Nisch and endogenous GLT-1 in comparison to GFP-NischΔphox mutant and GFP control. One-way ANOVA, post hoc Tukey’s test (*n* = 15-20 cells).

**Suppl. Figure 3:**

**A**) Confocal images of GFP-Nisch and GLT-1aBBS expressing astrocytes treated with 100μM glutamate for 0, 5, 30 and 60 min showed increased co-localization (shown in white) between GFP-Nisch (green) and GLT-1 compared to untreated controls.

**B**) The *Nisch* transgenic line was generated following the knockout-First strategy applied to the C57BL/6N Taconic strain. A L1L2_Bact_P cassette encoding an engrailed1 splice acceptor sequence, a LacZ reporter and a neomycin resistance gene was inserted between exons 4 and 5, disrupting *Nisch* transcription.

**C**) Xgal staining (dark blue) of sagittal brain section of a Nisch^HET^ animal demonstrates widespread Nischarin expression throughout the brain. Inset, hippocampus shows enriched Nischarin expression.

**D**) PCR analysis on DNA extracted from E16 WT, Nisch^KO^ and Nisch^HET^ embryos shows successful recognition of the WT and *Nisch* deletion alleles.

**Suppl. Figure 4:**

**A & B**) Astrocytes from WT and Nisch^KO^ hippocampal tissue cultures showed a similar membrane resistance (**A**), and resting potential (**B**).

## Materials and Methods

### Transgenic Animal

The *Nisch* (HEPD0811_2_A03; Allele: *Nisch*^*tm1a(EUCOMM)Hmgu*^) mouse line was obtained from the Wellcome Trust Sanger Institute as part of the International Knockout Mouse Consortium (IKMC) (Skarnes et al., 2011). The *Nisch* transgenic line was generated following the Knockout-First strategy on C57BL/6N Taconic strain. A L1L2_Bact_P cassette encoding an engrailed1 splice acceptor sequence, a LacZ reporter and a Neomycin resistance gene was inserted between exons 4 and 5 disrupting *Nisch* transcription. Animals were maintained under controlled conditions (temperature 20 ± 2°C; 12 hour light-dark cycle). Food and water were provided *ad libitum*. The genotyping was carried out following Sanger’s recommended procedures, briefly the DNA was extracted from ear biopsies and PCRs were performed with the following primers (5’ to 3’): Nisch_5arm_WTF: AGAGGCCCAGAGACCTGATA; Nisch_Crit_WTR: TGGACACGTGATGAGAAAGG; 5mut_R1: GAACTTCGGAATAGGAACTTCG; LacZ_2_small_F: ATCACGACGCGCTGTATC; LacZ_2_small_R: ACATCGGGCAAATAATATCG. All experimental procedures were carried out in accordance with institutional animal welfare guidelines and licensed by the UK Home Office in accordance with the Animals (Scientific Procedures) Act 1986.

### Yeast two-hybrid screen

This screen was done as described previously (Marie et al., 2002). Briefly, bait cDNA for the GLT-1 N terminus (amino acids 1–44 of the rat protein sequence) was cloned into the yeast expression vector pPC97 in frame with the GAL4 binding domain. It was screened against a random-primed cDNA library from seizure-stimulated adult rat hippocampus cloned in the yeast expression vector pPC86 in frame with the GAL4 activation domain. Interacting proteins were identified by colony selection on plates lacking leucine, tryptophan, and histidine and confirmed by using a β-galactosidase assay and by checking that in the absence of GLT-1 bait the library protein did not activate the reporter genes (*His3,* allowing growth on histidine-deficient medium, and *LacZ,* expressing β-galactosidase).

### Plasmid constructs

Mouse GLT-1a cDNA with V5 and HA epitope tag inserted into the extracellular loop of the transporter (between Pro^199^ and Pro^200^) and cloned into pcDNA3.1-was gifted by Dr. M. Rattray (Peacey et al., 2009). Bungarotoxin binding sequence (BBS) tag was introduced by PCR into GLT-1a and 1b between the two proline residues (P199 and P200) in the extracellular loop using the following primers (written 5’ to 3’): *ccctggagccctaccctgacCCATCTGAGGAGGCC*; *agctctcgtagtatctccaaGGTGCCACCAGAACTTT,* where lowercase text corresponds to the sequence of the BBS tag. Mouse Nischarin vector (clone ID: 100068156) was obtained from I.M.A.G.E. Consortium. The phox domain of Nischarin was deleted using the following primers: GTAAATGGTGTCACTGCAGCACT, CTCAGGGCCGAAGCTGAGTGT. The dominant negative, phox vector was generated using the following primers: GGCCTCATGGGCCCAG, TTCATAGAGGTGAAAATGCAGGA. The full length Nischarin, Dphox and phox vectors were C-terminally tagged with GFP by infusion cloning in frame into CAG-GFP (Addgene) using the following primers: ATCATTTTGGCAAAGCTAGCaccatggcggctgcgacact and CGTCGACTGCAGAATTCtgccagtgagctccacaggc, where lowercase text corresponds to parts of the Nischarin sequence. Neuronal processes were targeted using the RGECO vector obtained from Addgene.

### Preparation and transfection of astrocyte cultures

Primary cultures of cortical astrocytes were prepared from E18 or P0 Sprague-Dawley rats as previously described (Banker G, 1998). Cells were maintained in Dulbecco’s modified Eagle’s medium DMEM GlutaMAX (Invitrogen) supplemented with 4.5 g/L glucose, 20% fetal bovine serum, 10 u/ml penicillinG, and 100 μg/ml streptomycin at 37°C with 5% CO2 in a humidified incubator. Media was exchanged the day after plating. Astrocytes were passaged when confluency was reached (10 days after plating). For biotinylation assay, astrocytes were transfected with GLT-1aV5 and GFP (2μg, 1μg) or GLT-1aV5 and GFP-Nisch (2μg, 2μg) using Amaxa Nucleofector^®^ technology following the manufacturer’s protocol and maintained for 5 to 7 days before processing.

### Preparation and transfection of mixed culture and the neuron-astrocyte co-cultures

Hippocampal cultures were obtained from E18 rat embryos as described previously with some modifications (Arancibia-Carcamo et al., 2009). In order to enrich the culture with astrocytes, the neurons were kept 24h after plating in attachment medium (Minimal Essential Medium, 10% Horse Serum, 1 mM Sodium Pyruvate and 0.6% Glucose) before replacing with maintenance medium (Neurobasal Medium, B27 supplement, Glutamax, 0.6% Glucose, Penstrep). For optosplit experiments, after transfecting DIV7 hippocampal neurons with RGECO (lipofectamine 2000), astrocytes were transfected by nucleofection with GFP-Nisch (2μg) or GFP (1μg) (Amaxa Nucleofector) and plated on top of the RGECO transfected neurons. Transfected astrocytes were maintained with neurons for 3 to 4 days before multi-wavelength live-imaging. For live and fixed time lapse confocal and SIM imaging, astrocytes were transfected by nucleofection with GLT-1aBBS and GFP-Nisch (2μg, 2μg) or GLT-1aBBS and GFP (2μg, 1μg) (Amaxa Nucleofector) and plated on top of DIV10 hippocampal neurons. Transfected astrocytes were maintained with neurons for 3 to 4 days before imaging.

### Cell culture and transfection

COS7 and HeLa cells were maintained in 10cm dishes containing 10 ml Enhanced Dulbecco’s Modified Eagles Medium (DMEM), supplemented with pen/strep and 10% FBS, at 37°C and 5% CO2, transfected by nucleofection using an Amaxa electroporator and allowed 24-48 hours for protein expression.

### Coimmunoprecipitation assays from rat brain homogenate and cell culture

Coimmunoprecipitation experiments from brain/cell culture were performed as previously described (Twelvetrees et al., 2010). Briefly, mouse brain/cell culture expressing proteins of interest was homogenised in pull-down buffer (50 mM TRIS pH 7.5, 0.5 % triton X-100, 150 mM NaCl, 1 mM EDTA, 1mM PMSF with antipain, pepstatin and leupeptin at 10 μg/ml) and solubilised for 2 hours. Solubilised material was ultracentrifuged at 66,000 g for 40 minutes at 4°C and the supernatant (solubilised protein) was incubated with 2μg of anti-GLT1 (Alomone, Cat No: AGC022) antibody overnight at 4°C. To precipitate complexes, 20 μl protein-A or −G beads or GFP trap beads (Chrometek) were added for 1 hour at 4°C. Beads were then washed extensively and bound complexes were analysed by SDS-PAGE and western blotting.

### Western Blotting

SDS - polyacrylamide gel electrophoresis (PAGE) and Western Blotting samples were denatured at 94°C for 5 minutes in 3 x SDS sample buffer (150mM Tris pH 8, 6% SDS, 0.3M DTT, 0.3% Bromophenol Blue, 30% glycerol). Polyacrylamide gels were prepared using 10% running gels and 5% stacking gels in Novex 1.5mm Cassettes and run using the Novex XCell SureLock Mini-Cell system. Gels were transferred onto Hybond-C nitrocellulose membrane (GE Healthcare). Membranes were blocked in 4% milk for 1 hour and incubated overnight at 4°C with shaking in the appropriate primary antibody against Nischarin (1:500, BD Biosciences, Cat No: 558262), GFP (1:500, Santacruz, Cat No: sc-8334), V5 (1:2000, Invitrogen, Cat. No: R960-25), GLT (1:500, Alomone). HRP-conjugated secondary antibodies were from Rockland (1:10,000). Bands were visualised using Crescendo Chemiluminescent substrate (Millipore) together with an ImageQuant LAS 4000 CCD camera system (GE Healthcare).

### GST pull down assays from transfected COS7 cells

GST fusion with amino acids 1–44 of the N terminus of GLT-1 (GST–GLT-N), C terminus of GLT-1 (GST-GLT-C) and C terminus of GLAST (GST-GLAST-C), GST fusions A–E, encoding amino acids 1–15, 15–30, 30–44, 9–23, and 23–37 of GLT-1 were cloned as described previously (Marie et al., 2002). Pull-downs from brain were performed as described previously (Smith et al., 2008, 2010). Briefly, COS cells transfected with GFP-Nisch (2μg) was homogenized in pull-down buffer (50 mM HEPES, pH 7.5, 0.5% Triton X-100, 150 mM NaCl, 1 mM EDTA, and 1 mM PMSF with antipain, pepstatin, and leupeptin at 10g/ml) and solubilized for 2 h. Solubilized material was ultracentrifuged at 66,000g for 40 min at 4°C, and the supernatant (solubilized protein) was exposed to 10–20μg of GST fusion protein attached to glutathione–agarose beads for 1hr at 4°C. Beads were then washed extensively and analyzed by SDS-PAGE and western blotting.

### Biotinylation Assay

Surface biotinylation assays have been fully described previously (Smith et al., 2010; Twelvetrees et al., 2010). Briefly confluent astrocyte cultures were incubated on ice with biotin solution (Sulpho-NHS-biotin(PIERCE) at 0.5 mg/ml in PBS containing Ca^2+^ /Mg^2+^) and quenched with quench buffer (PBS Ca2+/Mg2+containing 1 mg/ml BSA). Astrocytes were solubilised for 1 hour in RIPA buffer (50 mM Tris pH 7.5, 1mM EDTA, 2 mM EGTA, 150 mM NaCl, 1% NP40, 0.5% DOC, 0.1% SDS, and 1 mM PMSF with antipain, pepstatin and leupeptin 10μg/ml) and the lysates were then centrifuged to pellet cell debris. 15% of the supernatant was taken to use as a total protein sample and the remainder was incubated for 2h with 25μl Ultralink immobilized NeutrAvidin (PIERCE) 50% slurry at 4 °C to precipitate biotin labeled membrane proteins. Beads were washed three times in RIPA buffer and analysed by SDS-PAGE and western blotting. Biotinylated surface GLT-1 transporters were identified by using anti-GLT primary antibody (1:500, Alomone) or anti-V5 primary antibody (1:2000, Invitrogen) and detection of enhanced chemilluminescence from HRP-coupled anti-rabbit secondary antibodies followed by detection with an ImageQuant LAS4000 mini imaging system and analysis with ImageQuant software (GE Healthcare).

### Excitotoxicity Assay

DIV14 hippocampal cultures were treated for 20 minutes with 10μM Glutamate and 10μM Glycine before replacement with conditioned maintenance media. 24 hours later, cultures were treated with 10μg/mL Propidium Iodide (PI) for 10 minutes prior to fixation for 5 minutes with 4% PFA at room temperature. Cell nuclei were stained with DAPI. Coverslips were imaged by confocal microscopy using a 10x objective (0.8x digital zoom, 4 averaging, 1024×1024, bit depth 8). Laser power and gain were kept consistent within and across all experiments. Images were analysed using ImageJ Cell Counter plugin, manually counting DAPI+ve cells and PI+ve cells in the same field of view. 3-4 field of views were taken per coverslip, 3-6 coverslips taken per embryo. N= 3 litter-matched embryos from separate preps. Experiments were performed blinded during image acquisition and analysis.

### Proximity Ligation Assay

The *in-situ* proximity ligation assay (PLA) was used according to the manufacturer’s instructions (Olink Bioscience). Neurons were fixed in 4% PFA/30% sucrose, blocked (10% horse serum, 0.5% BSA, and 0.2% Triton X-100, 10 min at room temperature), and incubated with primary antibodies (1:500, anti-GLT (gift from Dr. N.Danbolt) and anti-Nischarin (1:100, Sigma, Cat. No: HPA023189). For control PLA, single primary antibody was applied. Cells were washed in 1 × PBS and then incubated with secondary antibodies conjugated to oligonucleotides. Ligation and amplification reactions were conducted at 37°C, before mounting and visualization with confocal laser scanning microscope. Images were thresholded and number of puncta and DAPI stained nuclei were manually counted for each image using the *Metamorph software.* Experiments were performed blinded during image acquisition and analysis.

### Antibody Feeding

For receptor internalization and recycling assays, HeLa cells were transfected with GFPNisch and GLT-1a-HA (2μg, 2μg) or GFP and GLT-1a-HA (1μg, 2μg). The transporters were live labeled with anti-HA antibody in DMEM + 25mM HEPES at 17°C. Labelled transporters were allowed to internalize for 60 min at 37°C. For recycling assay, surface transporters were stripped using acid wash (0.2M acetic acid and 0.5M NaCl). Cells were then returned to the incubator for 30 min and 60 min at 37°C to allow internalized transporters to recycle to the surface. For surface staining, cells were fixed with 4% paraformaldehyde (PFA)/4% sucrose/PBS, pH 7, for 5 min and blocked with block solution (PBS, 10% horse serum, and 0.5% BSA) for 10 min, followed by Alexa Fluor-555-conjugated anti-mouse secondary antibody for 1hr (1:400; Invitrogen). For identifying internalized transporters, cells were subsequently permeabilized with block solution containing 0.2% Triton X-100 for 10min, followed by Alexa Fluor-555-conjugated anti-mouse secondary antibody (1:1000; Invitrogen). After extensive washing, coverslips were mounted on microscope slides using ProLong Gold antifade reagent (Invitrogen) and sealed with nail varnish. Images were attained using the confocal laser scanning microscope. Image analysis was performed using *ImageJ* software. For quantification of receptor internalization, red fluorescence that was not colocalized with cyan fluorescence (surface receptors) was calculated as the internalized transporter population or vice versa for calculating the surface transporter population in the recycling assay and normalized to total transporter population. Rate of internalization/recycling at different time points is measured as fold change relative to 0min.

### Immunostaining

Hippocampal cell culture was fixed with 4% PFA/4% sucrose/PBS, pH7, for 5min. The cells were permeabilized with block solution (PBS, 10% horse serum, and 0.5% BSA, 0.2% Triton) for 10min. The cells were incubated with primary antibodies including, anti-GLT1 (1:500, Alomone), anti-EEA1 (1:500, BD Bioscience, Cat. No: 610456), anti-Map2 (1:1000) for 1hr in block solution, followed by incubation with Alexa Fluor-488 or 555 or 647-conjugated secondary antibodies (1:1000, Invitrogen) for 1hr. After extensive washing, coverslips were mounted on microscope slides using ProLong Gold antifade reagent (Invitrogen) and sealed with nail varnish. Images were attained using Zeiss LSM 700 upright confocal microscope with an Apochromat 63× oil immersion lens with 1.4 numerical aperture. Images were digitally captured using ZEN software with excitation at 488nm for GFP and Alexa-Fluor 488, 555nm for Alexa-Fluor 555 and 633nm for Alexa-Fluor 647 and conjugated secondary antibodies. Pinholes were set to 1 Airy unit creating an optical slice of 0.8μm. For measuring colocalization, the colocalization plugin of the *Metamorph* software was used. Fluorescence intensity was measured using the *ImageJ* software.

### Live Imaging

Structured Illumination Microscopy was performed using a Zeiss Elyra PS.1 equipped with 488, 555 and 642 nm lasers. Images were acquired with 63×1.4 NA oil immersion objective using pco.edge sCMOS camera and Zen 2012 image analysis software. Typically, images were acquired with 51μm grating and 3 rotations by exciting fluorophores with 1-3% laser intensity and 120-150 ms exposure time. Post-acquisition, images were processed with Zen 2012 using the SIM reconstruction module with default settings and drift corrections between the channels were performed with respect to 100nm Tetraspec fluorescent microspheres (Molecular probes). To create kymographs image sequences were opened within ImageJ. Curved processes were straightened using the “straighten” macro and kymographs created by the “multiple kymograph” macro. Resultant kymographs show the process along the x-axis and time across the y-axis.

### Multi wavelength imaging and analysis (Optosplit)

An inverted Zeiss Axiovert 200 microscope (63x 1.4 NA oil objective), attached to an Evolve (EMCCD) camera (Photometrics), fitted with an image splitter (Optosplit II, Cairn Research), allowed simultaneous acquisition of images at two separate emission wavelengths. Videos were recorded at 8.5 Hz using Micro-manager software (Edelstein et al., 2010). Excitation was achieved through a D470/40X filter (Chroma) and emission was split using a 565DCXR dichroic beam-splitter (Chroma), subsequently collecting with HQ522/40M and HQ607/75M (Cairn Research) filters for GFP and RGECO, respectively. A Grass S9 stimulator and a stimulation bath (Warner Instruments) allowed field stimulation (10 Hz for 10 s) of neuron-astrocyte co-cultures prepared as described previously. Movies were aligned using the Cairn Image Splitter plugin in ImageJ. Graphs showing F/F_0_ were plotted using *Graph pad prism*. Regions of interest were manually drawn and fluorescence was normalized to the first 10 frames before stimulation.

### Image and statistical analysis

All experiments were performed on astrocytes/cell culture/mixed hippocampal astrocyte-neuro co-culture from at least three individual preparations. For all quantified experiments the experimenters were blind to the condition of the sample analyzed. All image analysis was performed blinded. Values are given as mean ± standard error of the mean (SEM). Error bars represent SEM. Statistical analysis was performed in GraphPad Prism (version 8; GraphPad Software, CA, USA) or Microsoft Excel. All data was tested for normal distribution with D’Agostino & Pearson test to determine the use of parametric (Student’s t test, one-way ANOVA) or non-parametric (Mann-Whitney, Kruskal-Wallis) tests. When p < 0.05, appropriate post hoc tests were carried out in analyses with multiple comparisons and are stated in the figure legends.

### Electrophysiology

Electrophysiology was carried out on embryonic hippocampal cultures on their tenth day *in vitro*. Cultures were prepared from nischarin WT or KO mice embryos at 16 days post-fertilization. Recordings were carried out at room temperature (20-21°C) with a HEPES-buffered extracellular solution mimicking cerebrospinal fluid (artificial cerebrospinal fluid, aCSF) containing (in mM): 140 NaCl, 10 HEPES, 10 glucose, 2.5 KCl, 2 CaCl_2_, 1 NaH_2_PO_4_, 1 MgCl_2_, pH 7.4 adjusted to with NaOH, osmolarity 300 mOsm (oxygenated with 100% O_2_). The solution was perfused at a flow rate of 3-4 ml/min through the recording chamber using gravity-driven perfusion from syringe barrels (60ml) connected to individual tubes, which merged into a single outlet just prior to reaching the bath. Whole-cell patch-clamp recordings were made from astrocytes using a potassium gluconate based internal solution, containing (in mM): 130 K-gluconate, 4 NaCl, 10 HEPES, 1 CaCl_2_, 10 EGTA, 2 MgATP, 0.5 Na_2_GTP (adjusted to pH 7.1–7.2 with KOH, and osmolarity ~ 285 mOsm). Alexa Fluor 594 (20 μM) was added to each aliquot of internal solution on the day of the experiment. Astrocytes were recognised visually by their low contrast soma, with an angular morphology and stellate processes revealed by dye-filling and usually gap junctional coupling allowing dye spread to other nearby astrocytes (Fig. 4C).

### Recording glutamate uptake in astrocytes

In order to record the glutamate uptake current from astrocytes, voltage-clamp recordings were made at the cell’s resting potential (typically around −90 mV). D-aspartate (200 μM, Sigma) was used to evoke a transporter current, since it is taken up by glial glutamate transporters (Davies & Johnston, 1976; Barbour *et al.*, 1991; Furness et al., 2008) and may have less effect than glutamate on glutamate receptors. However, D-aspartate may activate NMDA receptors or inhibit AMPA/KA receptors (Gong et al., 2005), or release glutamate via heteroexchange on transporters (Volterra et al., 1996). Any resulting activation of glutamate receptors might cause membrane potential depolarisation, neuronal action potentials and a rise of [K^+^]_o_ into the extracellular space, which could evoke an inward current in astrocytes (which have a highly K^+^-permeable membrane: Meeks & Mennerick, 2007). To prevent these effects we therefore supplemented the aCSF with a selection of blockers, which were present throughout the experiment: TTX to block action potentials (150nM, Tocris), a GABAA receptor blocker (bicuculline 10 μM, Sigma), NMDA receptor blockers (D-AP5 50 μM, Tocris; (+)MK-801 10 μM, Sigma; 5,7-DCK 10 μM, Sigma), an AMPA and kainate receptor blocker (NBQX 10 μM, Sigma), and an inwardly rectifying potassium channel blocker (barium chloride 200 μM, Sigma) which does not affect glutamate transport (Barbour et al., 1991). A non-transported glial glutamate transporter blocker (Shimamoto *et al*., 2004), TFB-TBOA (10 μM, Tocris) was also used in some experiments to block the glutamate transporter current evoked by D-aspartate. The size of the uptake current was calculated as the inward current recorded in D-aspartate minus the average of the baseline currents measured before and after D-aspartate application (using Clampfit 10.4). Experiments were performed with the experimenter blinded to the genotype.

