## Supplementary figures and images for "The non-adrenergic imidazoline-1 receptor protein Nischarin is a key regulator of astrocyte glutamate uptake"

### Supplemental figure 1

Supplementary Figure 1

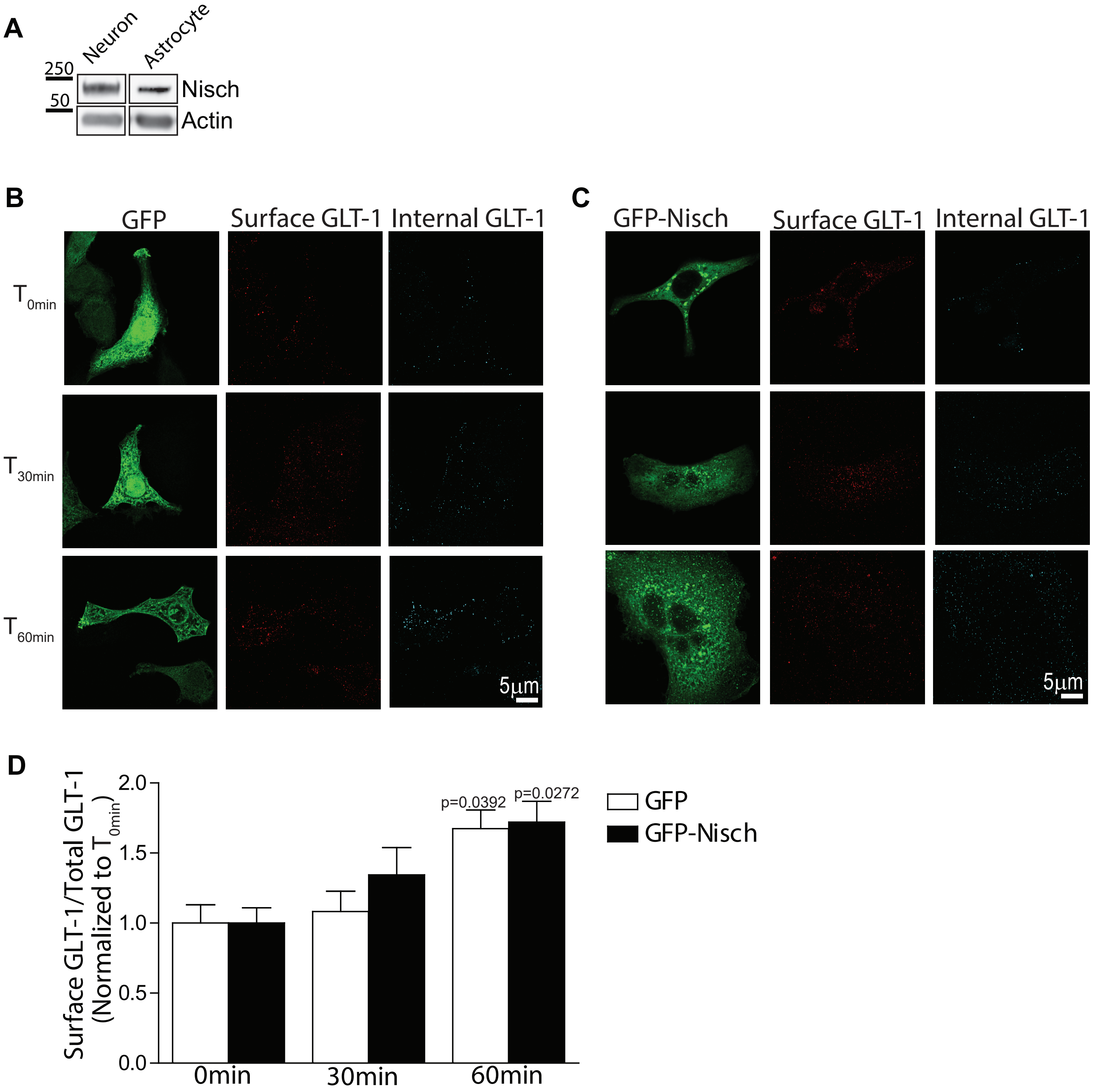

### Supplemental figure 2

Supplementary Figure 2

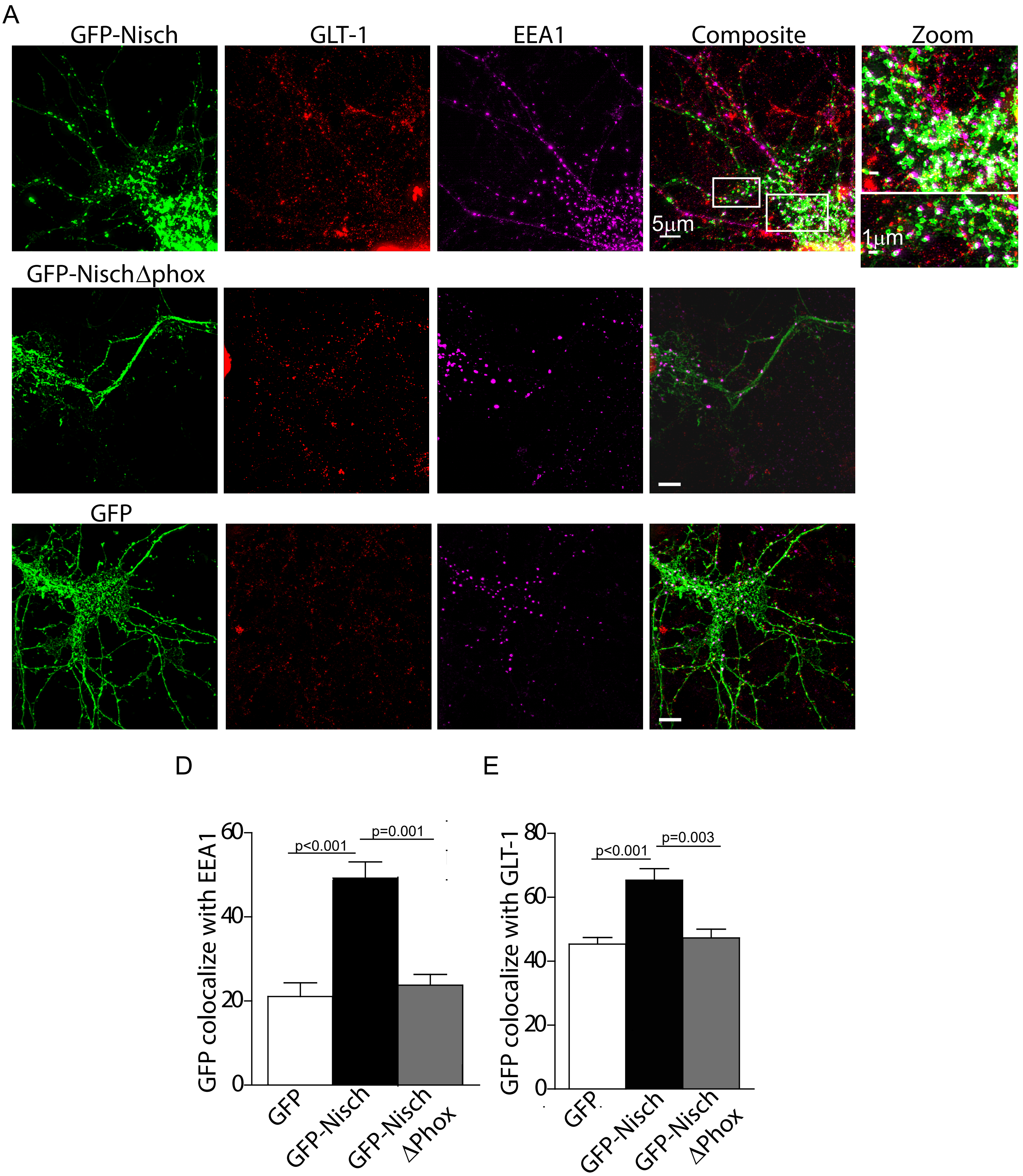

### Supplemental figure 3

Supplementary Figure 3

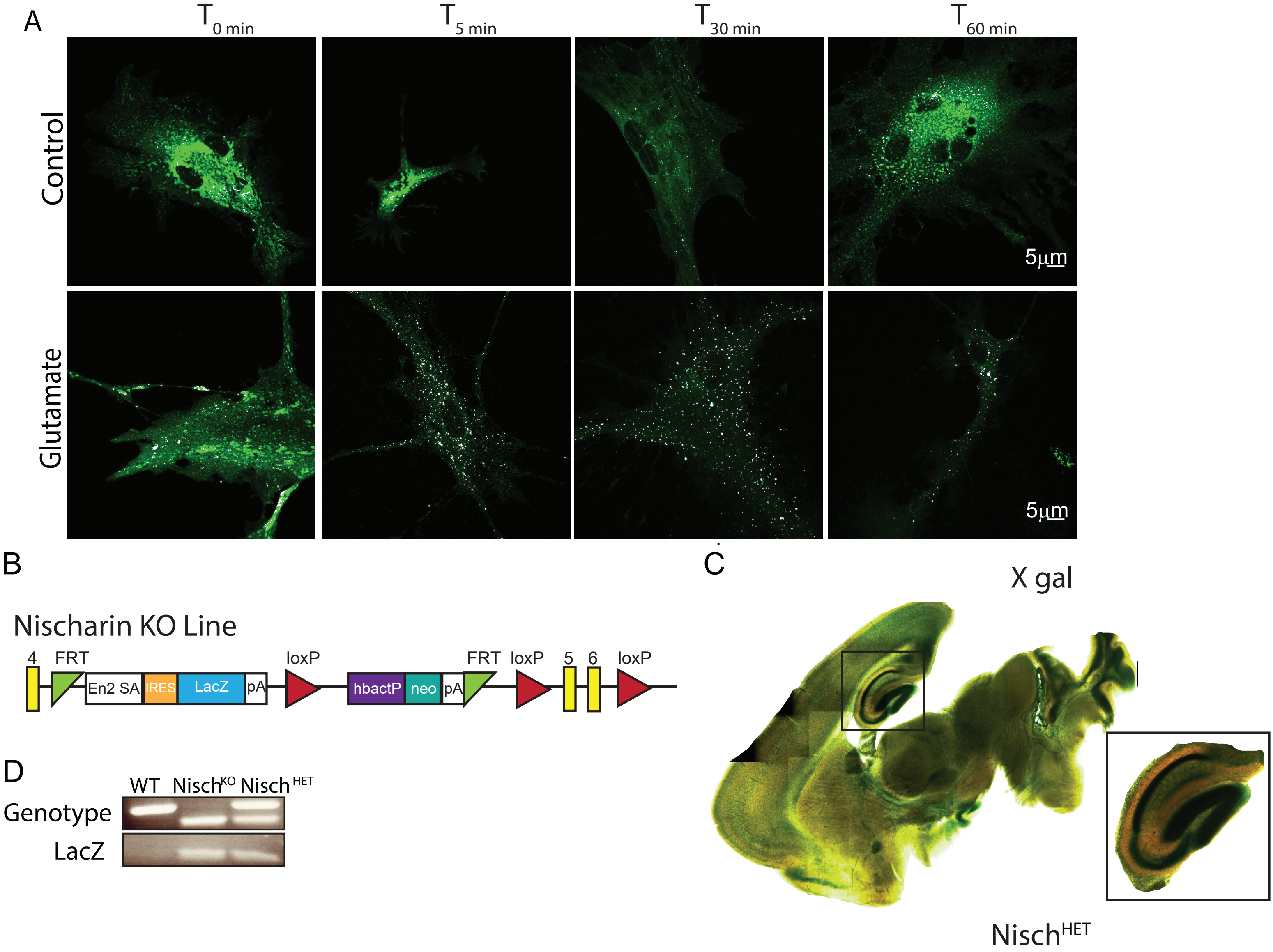

### Supplemental figure 4

Supplementary Figure 4

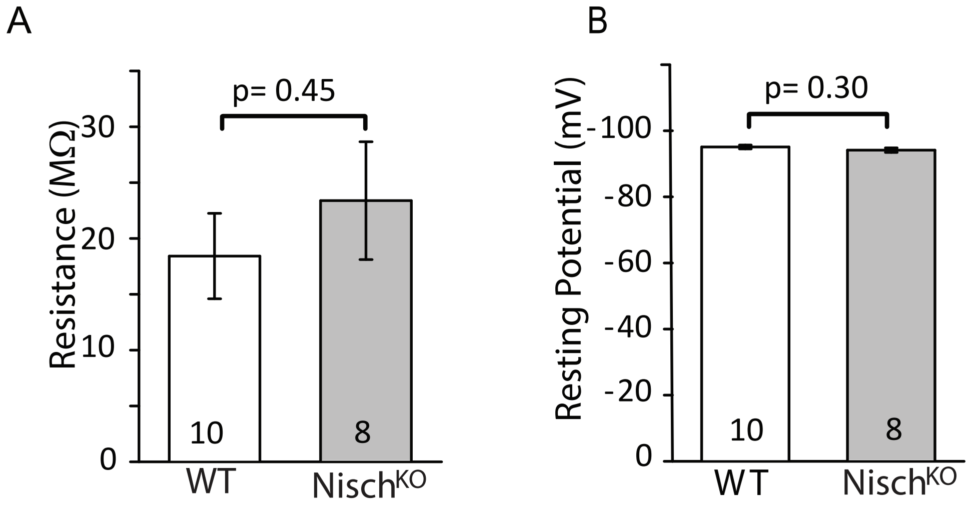
